# CoxMKF: A Knockoff Filter for High-Dimensional Mediation Analysis with a Survival Outcome in Epigenetic Studies

**DOI:** 10.1101/2022.06.29.498077

**Authors:** Peixin Tian, Minhao Yao, Tao Huang, Zhonghua Liu

## Abstract

**Motivation:** It is of scientific interest to identify DNA methylation CpG sites that might mediate the effect of an environmental exposure on a survival outcome in high-dimensional mediation analysis. However, there is a lack of powerful statistical methods that can provide a guarantee of false discovery rate (FDR) control in finite-sample settings.

**Results:** In this article, we propose a novel method called CoxMKF, which applies aggregation of multiple knockoffs to a Cox proportional hazards model for a survival outcome with high-dimensional mediators. The proposed CoxMKF can achieve FDR control even in finite-sample settings, which is particularly advantageous when the sample size is not large. Moreover, our proposed CoxMKF can overcome the randomness of the unstable model-X knockoffs. Our simulation results show that CoxMKF controls FDR well in finite samples. We further apply CoxMKF to a lung cancer data set from The Cancer Genome Atlas (TCGA) project with 754 subjects and 365 306 DNA methylation CpG sites, and identify four DNA methylation CpG sites that might mediate the effect of smoking on the overall survival among lung cancer patients.

**Availability:** The R package **CoxMKF** is publicly available at https://github.com/MinhaoYaooo/CoxMKF.

## 1 Introduction

Cigarette smoking is a well-known risk factor for inducing lung cancer (Walser et al., 2008). However, it’s unclear how epigenetic changes (e.g., DNA methylation) might mediate the effect of smoking on lung cancer survival. DNA methylation is the covalent bonding of a methyl group (CH_3_) onto the C5 position of the cytosine, which can regulate gene expressions by engaging the relevant genes or by blocking transcription factors binding to DNA (Moore et al., 2013). Since DNA methylation is a reversible process (Wu and Zhang, 2014), it’s thus of scientific interest to investigate the mediator role of DNA methylation CpG sites in the causal pathway from smoking to lung cancer survival for targeted intervention.

However, with high-dimensional DNA methylation data, it is challenging to select causal DNA methylation CpG sites with a guarantee of finite-sample false discovery rate (FDR) control. Recently, Luo et al. (2020) extended the Cox proportional hazards model (Cox, 1972) to high-dimensional mediators (e.g., CpG sites) settings by applying the minimax concave penalty (MCP) (Zhang, 2010) and Sobel test (Sobel, 1982) to select DNA methylation CpG sites as mediators. Furthermore, Zhang et al. (2021) introduced a de-biased Lasso estimator to handle ultra-dimensional mediators with a survival outcome. However, both methods cannot guarantee FDR control in finite-sample settings.

To address this challenge, we propose a novel mediator selection method named CoxMKF to identify DNA methylation CpG sites in the causal pathway from an environmental exposure to a survival outcome by incorporating aggregation of multiple knockoffs (AKO) with a guarantee of finite-sample FDR control (Nguyen et al., 2020). The key idea of the knockoffs inference is to generate fake copies of the original variables (conditionally on the original variables) and then perform variable selection (Candes et al., 2018). The model-X knockoff filter proposed by Candes et al. (2018) generates knockoffs by mimicking the original design matrix which are flagged as false positives in a regression model. The finite-sample FDR control is achieved by using a data-dependent threshold. Nonetheless, model-X knockoffs are generated by Monte Carlo sampling only once, resulting in instability due to randomness and difficulties in reproducing the study findings (Gimenez and Zou, 2019; Nguyen et al., 2020). AKO overcomes this limitation by repeating the model-X knockoffs generation procedure multiple times and then aggregates them together to increase stability (Meinshausen et al., 2009).

Our proposed CoxMKF method brings the idea of statistical aggregation from Meinshausen et al. (2009) to conduct variable selection via AKO (Nguyen et al., 2020) to improve stability. Specifically, our method has three steps. First, we take the *p*-values from the linear regression model for the exposure-mediator relationship, and then carry out the Benjamini-Hochberg (BH) procedure (Benjamini and Hochberg, 1995) to select top *d* significant DNA methylation CpG sites. Second, we generate model-X knockoffs according to the correlation structure of the *d* DNA methylation CpG sites. We then apply the Cox proportional hazards model (Cox, 1972) to the *d* DNA methylation CpG sites and their knockoff variables to obtain the regression coefficients. Third, we repeat the aforementioned steps for *B* times and propose two new feature statistics named Cox Coefficient Difference (CCD) and Multiple Cox Statistic (MCS) to determine a data-dependent threshold by applying statistical quantile aggregation with a pre-specified FDR level for variable selection. Our simulations indicate that the proposed CoxMKF may effectively control FDR with better stability in finite-sample settings. We further apply CoxMKF to a lung cancer data set from The Cancer Genome Atlas (TCGA) project and finally identify four DNA methylation CpG sites that might mediate the effect of smoking on lung cancer survival.

The rest of this article is organized as follows. In Section 2, we describe the regression models and introduce our CoxMKF procedure for high-dimensional mediation analysis with a survival outcome. In Section 3, we evaluate the performance of CoxMKF across a wide range of simulation settings. In Section 4, we apply CoxMKF to the lung cancer data from the TCGA project. Section 5 provides some concluding remarks.

## 2 Methods

Let *D_i_* be the duration of an event (e.g., lifespan) of individual *i*, and *C_i_* be the corresponding censoring time. The observed survival outcome is *T_i_* = min{*C_i_*, *D_i_*}, and the censoring indicator is *δ_i_* = *I*(*D_i_* ≤ *C_i_*). We evaluate the causal mechanism from an exposure *X_i_* to a survival outcome *T_i_* through a *p*-dimensional mediator vector ***M***_*i*_ = (*M*_*i*1_, … , *M_ip_*)^⊤^ (see Figure **??**). If *p* = 1, this problem reduces to a single mediator setting. In the multiple mediators setting (*p* ≥ 2), we perform mediation analysis using the following two regression models:

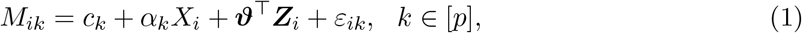

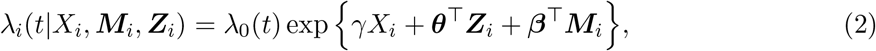

where [*p*] denotes the set of {1, 2, … , *p*}, *λ_i_*(*t*|*X_i_, **M**_i_, **Z**_i_*) is the Cox proportional hazards model, *λ*_0_(*t*) is an unspecified baseline hazard function, ***Z***_*i*_ = (*Z*_*i*1_, … , *Z_iq_*) is a *q*-dimensional vector of the baseline covariates (e.g., gender, age), and *ε_ik_* is the noise term. In model (1), ***α*** = (*α*_1_, … , *α_p_*)^⊤^ is a parameter vector relating the exposure to the mediators, and ***c*** = (*c*_1_, … , *c_p_*)^⊤^ is the intercept term. In model (2), *γ* is the direct effect of the exposure on the outcome, and ***β*** = (*β*_1_, … , *β_p_*)^⊤^ represents the effect of the mediators on the survival outcome adjusting for the effect of the exposure. ***ϑ*** and ***θ*** are the regression coefficients of the covariates on the mediators and on the survival outcome, respectively.

We use the potential outcomes framework (Rubin, 1974; Splawa-Neyman et al., 1990) to decompose the effect of the exposure on the outcome. For the Cox proportional hazards model, we can decompose the effect using the difference in counterfactual log hazards (VanderWeele, 2011; Huang and Yang, 2017). Specifically, let *T* (*x, **m***) denotes the survival time when the exposure is set to *x* and the vector of mediators is set to ***m***, and ***M*** (*x*) = (*M*_1_(*x*), … , *M_p_*(*x*))^⊤^ denotes the vector of mediators when the exposure is set to *x*. Let ***Z*** denotes the baseline covariates, and *x*\* denotes the reference level of the exposure. The total effect (TE) of the exposure on the outcome is defined as log *λ*(*T* (*x*, ***M*** (*x*)); *t*|***Z***) − log *λ*(*T* (*x*\*, ***M*** (*x*\*)); *t*|***Z***); the natural direct effect (NDE) is defined as log *λ*(*T* (*x*, ***M*** (*x*\*)); *t*|***Z***)−log *λ*(*T* (*x*\*, ***M*** (*x*\*)); *t*|***Z***); and the natural indirect effect (NIE) is defined as log *λ*(*T* (*x*, ***M*** (*x*)); *t*|***Z***) − log *λ*(*T* (*x*, ***M*** (*x*\*)); *t*|***Z***).

The following equation summarizes the relationship among these effects:

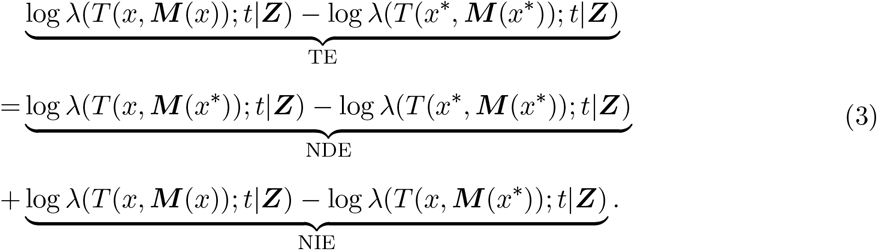

According to VanderWeele (2011) and Huang and Yang (2017), the following standard assumptions are required to ensure the identification of NDE and NIE:

(C1) *T* (*x, **m***) ⊥ *X*|***Z***: no unmeasured confounding between the exposure and the outcome.
(C2) *T* (*x, **m***) ⊥ *M_k_*|*X*, ***Z*** for *k* ∈ [*p*]: no unmeasured confounding between mediators and the outcome.
(C3) *X* ⊥ *M_k_*|***Z*** for *k* ∈ [*p*]: no unmeasured confounding between the exposure and mediators.
(C4) *T* (*x, **m***) ⊥ *M_k_*(*x*\*)|***Z*** for *k* ∈ [*p*]: no exposure-induced confounding between the mediators and the survival outcome.

If assumptions (C1) - (C4) hold, then the counterfactual log hazard can be approximated as:

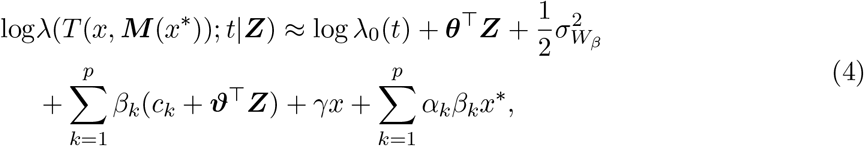

where 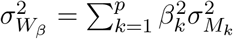 and 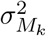 is the variance of the *k*th mediator (Huang and Yang, 2017; Luo et al., 2020). Therefore, both NDE and NIE can be estimated on the log hazard scale. In our models (1) and (2), we further specify the expressions of NDE and NIE in equation (3):

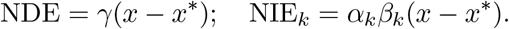

One main challenge here is that when the dimension *p* is much larger than the sample size *n*, the Cox proportional hazards model (2) cannot be fitted directly. To address this challenge, we propose a novel framework for variable selection of the mediators that can provide finite-sample FDR control, which contains three major steps (as shown in Figure 2).

**Figure 1:**
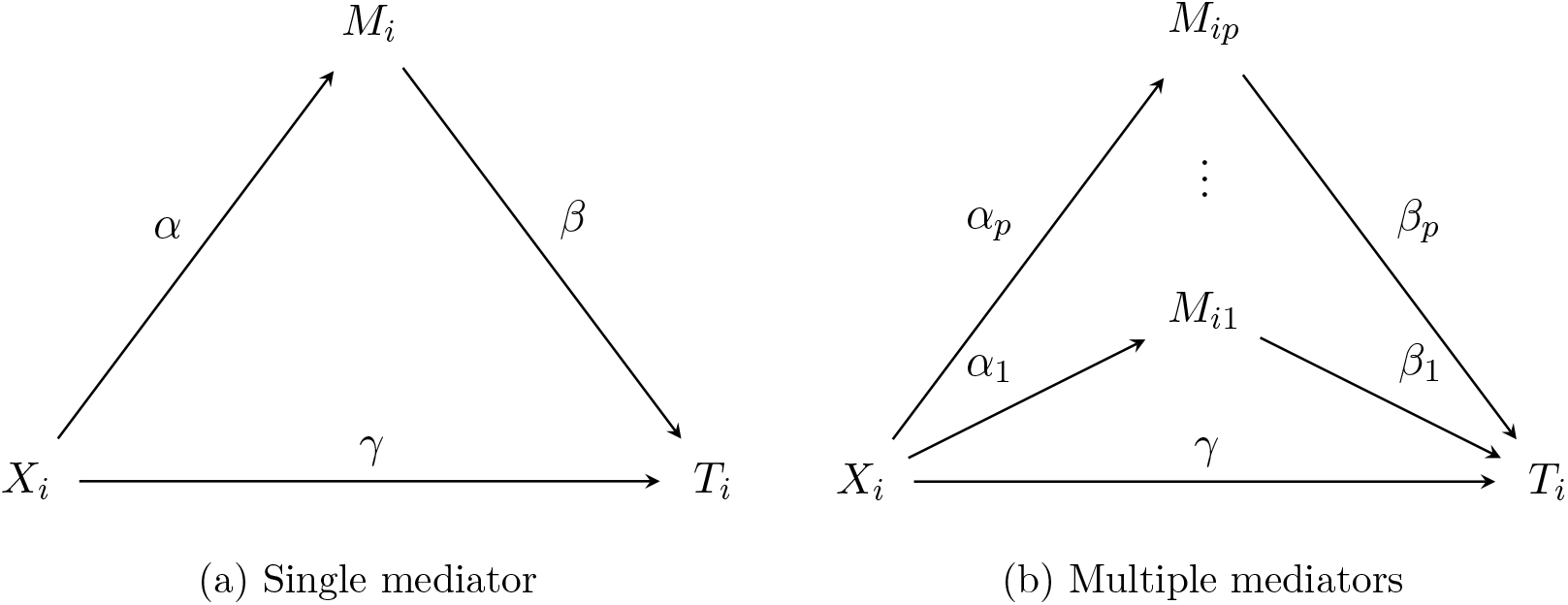
Causal diagram. The single mediator(left) and multiple mediators(right) related with the exposure variable *X_i_* and the outcome *T_i_*.

**Figure 2:**
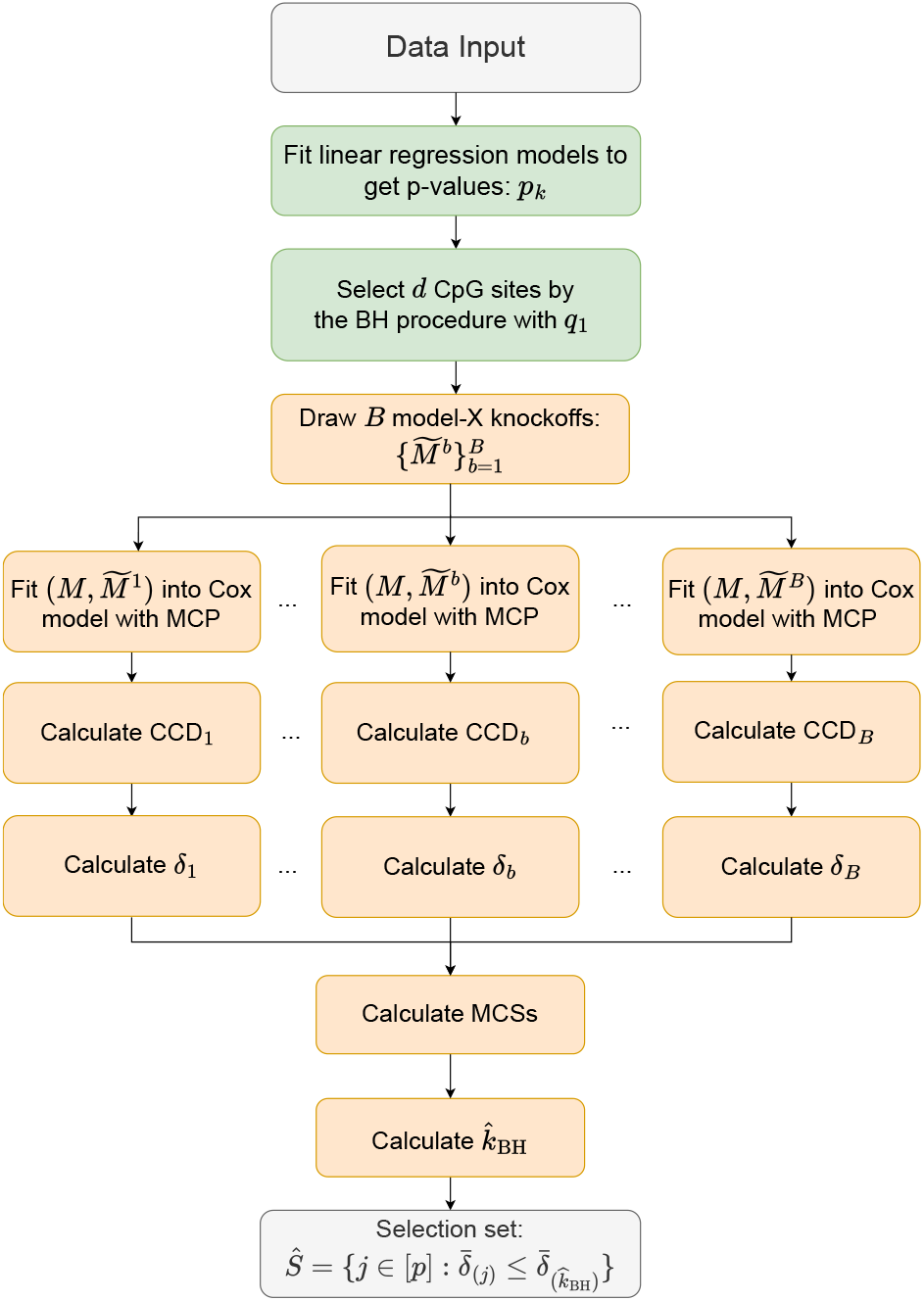
Flowchart of CoxMKF. First, we perform variable screening using the BH procedure in the *X_i_* → *M_ik_* relationship to obtain *d* mediators. Second, we sample *B* model-X knockoffs to fit in the Cox model. Third, we compute CCDs and MCSs to select the candidate mediators with a pre-specified FDR level.

First, we apply variable screening to the mediators using the Benjamini-Hochberg (BH) procedure (Benjamini and Hochberg, 1995). Specifically, we obtain the *p*-value in the *X_i_* → *M_ik_* relationship and select a subset of significant mediators of size *d* by controlling the FDR at level *q*_1_.

Second, we introduce aggregation of multiple knockoffs (AKO) (Nguyen et al., 2020) to control the finite-sample FDR in the selection of mediators with non-zero NIEs on the survival outcome. Suppose that we have *n* individuals. We denote ***M*** = (***M***_1_, … , ***M**_n_*)^⊤^ as the *n* × *d* design matrix of mediators after variable screening, and ***T*** = (*T*_1_, … , *T_n_*)^⊤^ as the vector of the observed survival outcomes. We generate *B* model-X knockoffs 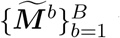 using the R package ‘knockoff’ (Candes et al., 2018). Moreover, our procedure maintains two key properties of model-X knockoffs (also shown in Figure 3) (Candes et al., 2018; Nguyen et al., 2020):

1. For any subset *S* ⊆ [*d*] and *b* ∈ [*B*], 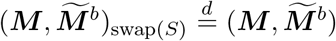, where 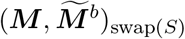 is obtained by swapping the *k*th columns of ***M*** and 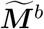 for each *k* ∈ *S*;
2. 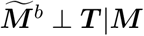 for any *b* ∈ [*B*].

**Figure 3:**
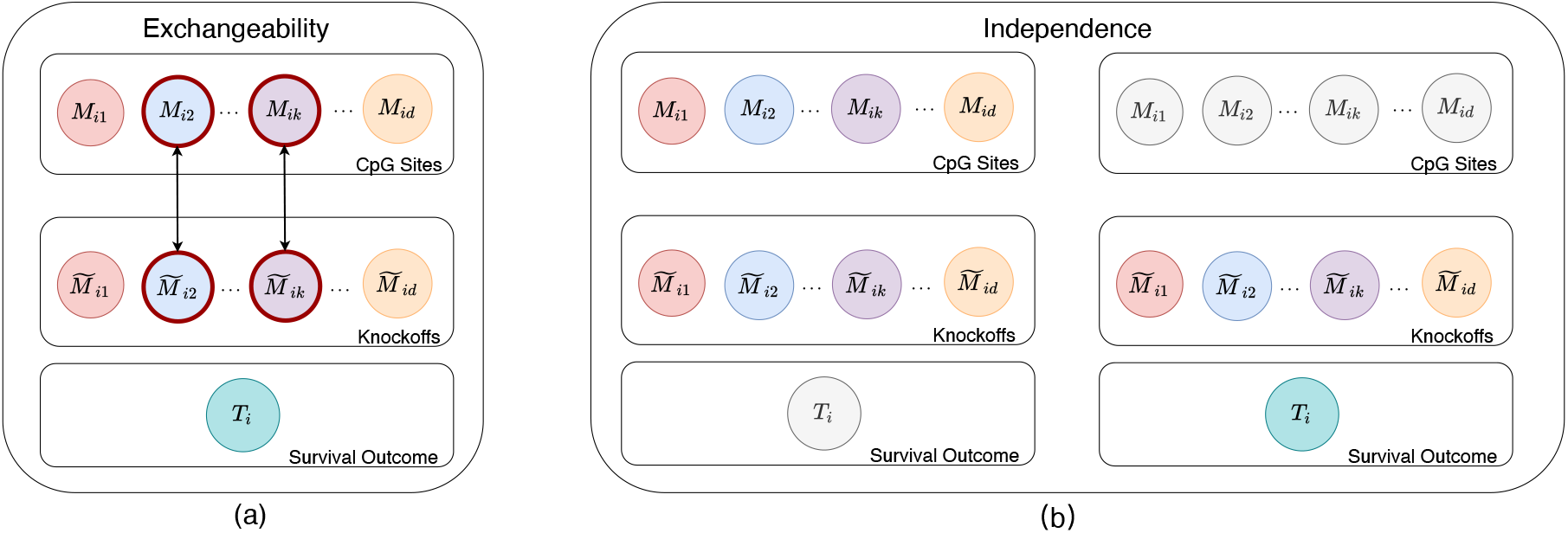
Illustration of knockoffs variables. (a) The knockoff copy keeps exchangeable property between ***M*** and 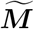. As the left penal shows, the knockoff variable 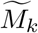 can exchange the position with *M_j_*. In this case, the model result will be unchanged if the positions of *M_k_* and 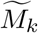 are swapped. (b) The knockoff copy shows independence with the response *T* (uncolored) when conditional on the original features (colored), and shows dependence with *T* (colored) when not conditional on the original features (uncolored).

It is worth noting that the second property is guaranteed if 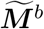 is constructed without looking at the response variable ***T***.

Third, we introduce a new aggregated feature statistic, called Multiple Cox Statistic (MCS), to select mediators with non-zero NIEs on the outcome. To calculate the MCS, we first obtain a *B*-by-*d* Cox Coefficient Difference (CCD) 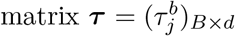 whose entries are:

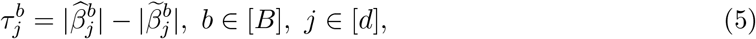

where 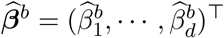 and 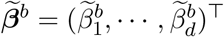 are the coefficients obtained by fitting the Cox proportional hazards model with the MCP using both the mediators ***M*** and the *b*th knockoffs 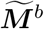. Based on the CCD matrix ***τ***, we then calculate another *B*-by-*d* matrix of intermediate statistics 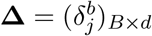 whose entries are:

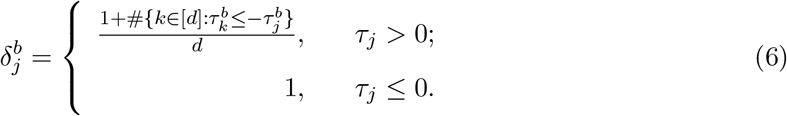

Generally, a small 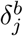 value suggests a strong non-null signal for the *j*th mediator in the *b*th replication. Following Meinshausen et al. (2009), we aggregate the intermediate statistics on *B* rows to generate the MCS for each candidate mediator respectively:

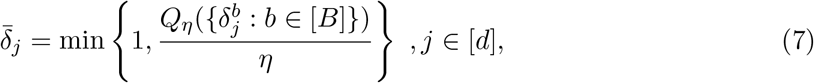

where *η* is the pre-specified quantile point, and *Q_η_*(·) denotes the *η*-quantile function. From the simulation, we find that our proposed CoxMKF becomes more conservative as *η* increases (as shown in Supplementary Figure S1). Therefore, we suggest using *η* = 0.05. Specifically, a small value of 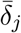 suggests that the *j*th mediator is considered to have a non-zero mediation effect over replications, and thus is more likely to be the true mediator. Following Nguyen et al. (2020). We then apply the BH procedure to determine a data-dependent threshold 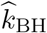 to control the FDR:

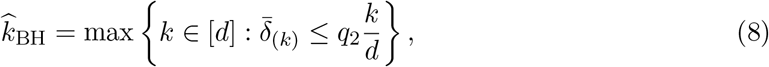

where *q*_2_ is a pre-specified target FDR level, and 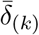 is the *k*th smallest value of 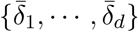.

Then we define the selection set of mediators with non-zero mediation effects as:

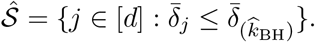

In this article, we evaluate the performance of CoxMKF by FDR and true positive proportion (TPP):

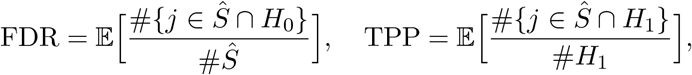

where *H*_0_ = {*k* : NIE_*k*_ = 0} includes the candidate mediators with no signal, and *H*_1_ = {*k* : NIE_*k*_ ≠ 0} includes the true mediators.

## 3 Simulation Studies

In this section, we conduct a wide range of simulation studies to evaluate the finite-sample performance of our proposed method. First, we generate the survival outcome ***T*** = (*T*_1_, *T*_2_, … , *T_n_*)^⊤^ from the exponential model *λ_i_*(*t*|*X_i_*, *Z*_*i*1_, *Z*_*i*2_, ***M**_i_*) = 0.5 exp {*γX_i_* + ***θ***^⊤^ ***Z***_*i*_ + ***β***^⊤^ ***M***_*i*_}, where the exposure *X_i_* is generated from a Bernoulli distribution with success probability 0.6 as Ber(0.6), the binary covariate ***Z***_1_ = (*Z*_1,1_, … , *Z*_*n*,1_)^⊤^ and the continuous covariate ***Z***_2_ = (*Z*_1,2_, … , *Z*_*n*,2_)^⊤^ are generated from a Bernoulli distribution with success probability 0.3 as Ber(0.3) and a uniform distribution as *U* (0, 1), respectively, and the coefficients of the covariates on the outcome are set to be *θ*_1_ = 0.3 and *θ*_2_ = −0.2. Then we generate the mediators ***M*** = (*M_ik_*)_*n*×*p*_ according to model (1), where the coefficient *c_k_* is generated from a uniform distribution as *U* (0, 1), the noise *ε_ik_* is generated from a standard normal distribution as *N* (0, 1), and ***θ*** is fixed to be (0.3, 0.2)^⊤^. As for ***α*** and ***β***, we fix the first ten elements of ***α*** to be *κ* · (0.55, 0.45, −0.40, −0.45, 0.50, 0.60, −0.40, −0.46, −0.40, 0.50)^⊤^, and the first twelve elements of ***β*** to be *κ* · (0.52, 0.45, 0.40, 0.40, −0.54, −0.60, −0.40, −0.50, 0, 0, 0.40, −0.80), where *κ* is a scaling factor to avoid the TPP to be too close to 1. For *n* = 300, 500 and 800, we set *κ* = 1, 0.8 and 0.6, respectively. The rest elements of ***α*** and ***β*** are set to be 0. Thus, (*M_i_*_1_, *M_i_*_2_, … *M_i_*_8_) have non-zero causal mediation effects, and *H*_1_ = {1, 2, … , 8} is the set including these eight true mediators. The censoring time is generated from an exponential distribution with parameter *c*_0_. By denoting proper *c*_0_, the average censoring rates are controlled at 20%, 40% and 60%, respectively. All the scenarios are based on 500 replications, where the sample sizes *n* = (300, 500, 800) and the numbers of candidate mediators *p* = (500, 1 000, 10 000), respectively. We set *q*_1_ = 0.2 in the screening step, the targeted FDR level *q*_2_ = 0.05 in equation (8), and the quantile point *η* = 0.05 in equation (7). In addition, based on the recommendation of Nguyen et al. (2020), we set the number of bootstrap replications *B* to 25, which could empirically offer stable performance.

We compare CoxMKF to Luo et al. (2020)’s method (denoted as HIMAsurvival). We compute the FDR and TPP to assess the accuracy of variable selection, and the results are reported in Table 1 (the standard errors of FDR and TPP can be found in Supplementary Table S1). When the number of candidate mediators *p* is small, CoxMKF and HIMAsurvival can both control the FDR under the pre-specified level. However, the FDR of HIMAsurvival is inflated when *p* = 10 000 and *n* = 300 or 500. In addition, nearly half of FDR values of HIMAsurvival across 500 replications are higher than 0.05 when *n* = 300, *p* = 10 000 and the censoring rate is controlled at 60% (as shown in Figure 4). When the sample size *n* increases to 800, HIMAsurvival can almost achieve the FDR control with a slightly inflated FDR for *p* = 10 000. Overall, HIMAsurvival fails to control the finite-sample FDR when *p* increases to 10 000 and the sample size is not large. By comparison, CoxMKF can guarantee the FDR under the pre-specified level in all scenarios, even if the sample size is small and the number of candidate mediators is large. For example, when *n* = 300, *p* = 10 000 and the censoring rate CR= 60%, the FDR of HIMAsurvival and CoxMKF are 0.129 (> 0.05) and 0.033 (< 0.05) respectively, indicating a successful finite-sample FDR control of CoxMKF with slight conservativeness while HIMAsurvival cannot control FDR in this setting. We also conduct simulations with different values of ***α***, ***β*** and a different choice of parameters (*q*_1_ = 0.05, *q*_2_ = 0.02, *η* = 0.3, *B* = 50), and the results are shown in Supplementary Table S2 and S3. We find that the FDR of our proposed CoxMKF is still below the nominal level, while HIMAsurvival fails to control the FDR when *p* = 10 000 in the new simulation setup. Due to the inflated FDR of HIMAsurvival, the TPP of HIMAsurvival is generally higher than CoxMKF.

**Table 1:** The FDR and TPP of mediator selection across 500 simulations with *q*_1_ = 0.2, *q*_2_ = 0.05, *η* = 0.05 and *B* = 25. The number of true mediators is 8. CR stands for the censoring rate. *κ* is a scaling factor to shrink the coefficients so that TPP is not too close to 1.

| n | CR | p | FDR ($< 0.05$) | | TPP | |
| --- | --- | --- | --- | --- | --- | --- |
|  |  |  | CoxMKF | HIMAsurvival | CoxMKF | HIMAsurvival |
| 300 ($\kappa = 1.0$) | 20% | 500 | 0.025 | 0.010 | 0.826 | 0.879 |
|  |  | 1000 | 0.033 | 0.010 | 0.766 | 0.877 |
|  |  | 10000 | 0.022 | 0.091 | 0.554 | 0.878 |
|  | 40% | 500 | 0.039 | 0.013 | 0.797 | 0.828 |
|  |  | 1000 | 0.025 | 0.010 | 0.761 | 0.832 |
|  |  | 10000 | 0.029 | 0.086 | 0.548 | 0.839 |
|  | 60% | 500 | 0.040 | 0.007 | 0.781 | 0.765 |
|  |  | 1000 | 0.026 | 0.009 | 0.723 | 0.756 |
|  |  | 10000 | 0.033 | 0.129 | 0.537 | 0.763 |
| 500 ($\kappa = 0.8$) | 20% | 500 | 0.021 | 0.013 | 0.879 | 0.947 |
|  |  | 1000 | 0.019 | 0.017 | 0.812 | 0.932 |
|  |  | 10000 | 0.022 | 0.061 | 0.612 | 0.903 |
|  | 40% | 500 | 0.020 | 0.010 | 0.852 | 0.925 |
|  |  | 1000 | 0.026 | 0.014 | 0.809 | 0.918 |
|  |  | 10000 | 0.018 | 0.058 | 0.611 | 0.901 |
|  | 60% | 500 | 0.032 | 0.010 | 0.856 | 0.861 |
|  |  | 1000 | 0.028 | 0.013 | 0.805 | 0.853 |
|  |  | 10000 | 0.022 | 0.058 | 0.618 | 0.833 |
| 800 ($\kappa = 0.6$) | 20% | 500 | 0.015 | 0.011 | 0.837 | 0.927 |
|  |  | 1000 | 0.021 | 0.012 | 0.766 | 0.916 |
|  |  | 10000 | 0.020 | 0.051 | 0.527 | 0.916 |
|  | 40% | 500 | 0.015 | 0.009 | 0.822 | 0.912 |
|  |  | 1000 | 0.016 | 0.008 | 0.787 | 0.921 |
|  |  | 10000 | 0.020 | 0.052 | 0.550 | 0.905 |
|  | 60% | 500 | 0.034 | 0.010 | 0.802 | 0.796 |
|  |  | 1000 | 0.024 | 0.009 | 0.749 | 0.791 |
|  |  | 10000 | 0.023 | 0.054 | 0.511 | 0.807 |

**Figure 4:**
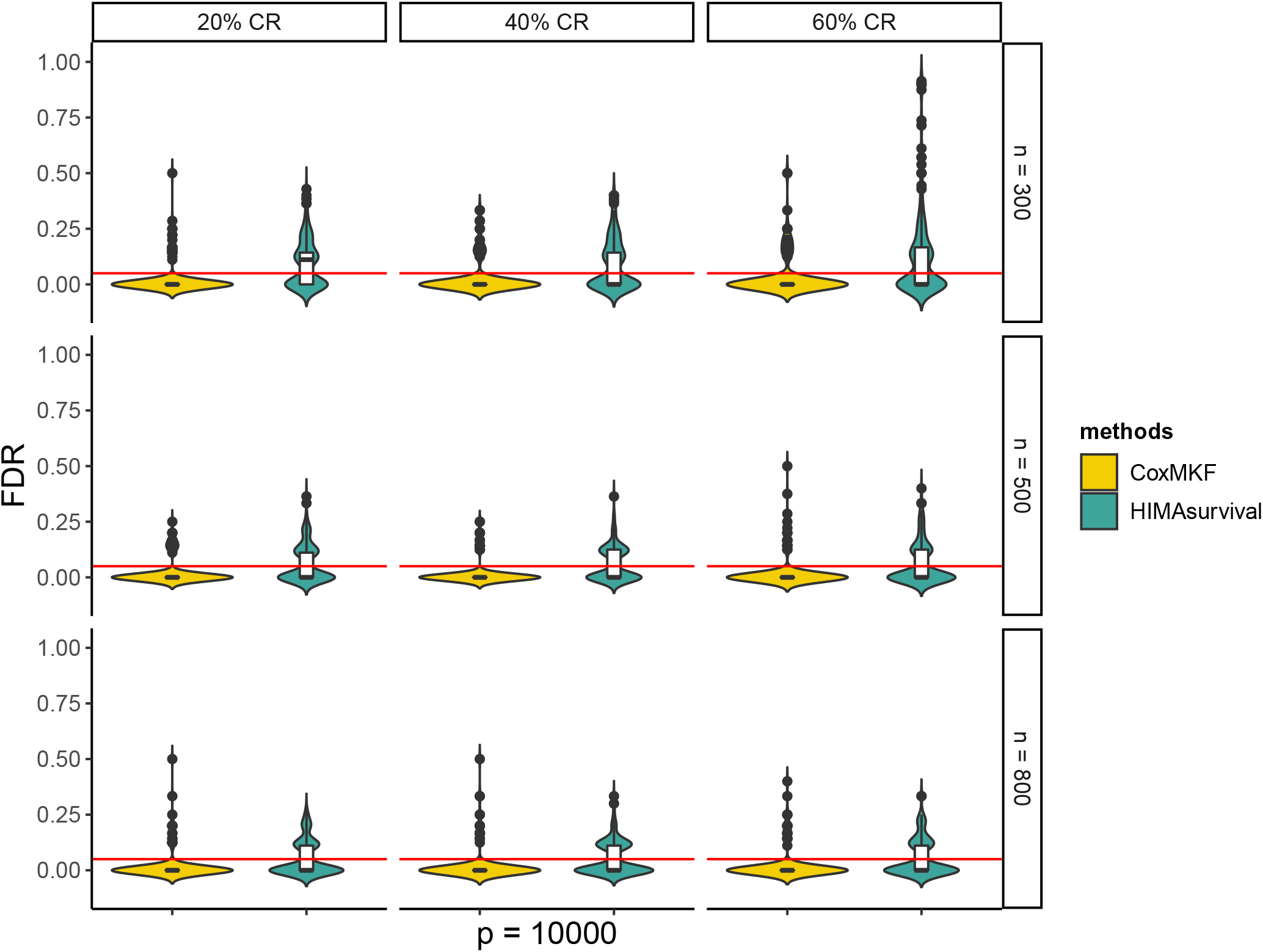
Violin plot and box plot of the FDR of CoxMKF and HIMAsurvival with *q*_1_ = 0.2, *q*_2_ = 0.05, *η* = 0.05, *p* = 10 000 and the number of AKO’s bootstrap replications *B* = 25 across 500 replications in simulations. CR stands for the censoring rate, and the red line represents the FDR control level *q*_2_ = 0.05. The number of true mediators is 8. We report the FDR when *p* = 10 000, and differentiate the methods by color, with yellow representing our proposed method CoxMKF and green representing the competing method HIMAsurvival. The width of the plot represents the value density. The greater the width of the shape, the denser the distribution of points.

To mimic the real data example, we also conduct a simulation with *n* = 800, *p* = 350 000 and the censoring rate CR= 60% over 500 replications. We set *q*_1_ = 5 × 10^−4^ in the preliminary screening, *q*_2_ = 0.1 as the targeted FDR level, *η* = 0.05 as the quantile point and the number of bootstrap replications *B* = 25.The FDR and TPP of CoxMKF are 0.006 and 0.652, respectively. By comparison, the FDR and TPP of HIMAsurvival are 0.146 (> 0.1) and 0.991, respectively. CoxMKF can therefore guarantee the finite-sample FDR control in high-dimensional settings. By comparison, HIMAsurvival cannot provide a guarantee of finite-sample FDR control in this case.

The simulation studies demonstrate that CoxMKF can provide a guarantee of finite-sample FDR control even when the censoring rate is high and the number of candidate mediators is large.

## 4 An Application to the Lung Cancer Data

Lung cancer is the most common cancer among men and the third most common cancer among women, and remains the leading cause of mortality (Sung et al., 2021). Among all the lung cancer diagnoses, approximately 85% of all diagnoses are non-small cell lung cancer (NSCLC) and 15% of them are small-cell lung cancer (Cao et al., 2018). There are many urgent challenges for researchers, such as finding an effective therapy and diagnosing the disease at early stage, which can significantly improve the treatment success rate and survival rate (Moore et al., 2013). Tobacco smoking is an important risk factor for lung cancer, and smoking status might affect the tumor genomic landscape (Govindan et al., 2012). Additionally, tobacco smoking is also associated with DNA methylation (Breitling et al., 2011). Therefore, it is of great interest to find DNA methylation CpG sites that mediate the effect of tobacco smoking on the lung cancer survival.

We apply our proposed CoxMKF to the lung cancer data from the TCGA project including lung squamous cell carcinoma and lung adenocarcinoma (https://xenabrowser.net/datapages/) to identify potential DNA methylation CpG sites between smoking and survival of lung cancer patients. In total, there are 754 patients aged 33-90 years with 365 306 DNA methylation CpG cites after excluding the patients with no survival time, smoking status and other covariates. The patients had DNA methylation profiles measured by the Illumina Infinium HumanMethylation 450 platform, and the DNA methylation values were recorded via BeadStudio software.

We are interested in detecting the DNA methylation CpG sites between smoking status (smoker/non-smoker) and lung cancer patients’ survival. The survival of lung patients is encoded as the number of days from the initial diagnosis to the death or the censoring time. The median survival time is 1 632 days, and we observe 305 deaths over the follow-up period with a censoring rate of 60%. We also include the following covariates: sex, age, cancer stage and radiotherapy indicator.

For CoxMKF, we set *q*_1_ = 5 × 10^−4^ in the screening step, *q*_2_ = 0.1 as the targeted FDR level, *η* = 0.05 as the quantile point and the number of AKO’s bootstrap replications *B* = 25. CoxMKF identifies four DNA methylation CpG sites (cg21926276, cg24129177, cg24200525 and cg07690349) in total, and the results of the effect estimates and the corresponding *p*-values are reported in Table 2. Among the three DNA methylation CpG sites, cg21926276 and cg24129177 have positive NIEs, through which smoking increases the possibility of mortality. The first methylation biomarker, cg21926276, is located in gene H19 on chromosome 11, and the estimated NIE is 0.149. The CpG site cg24129177 is located on chromosome 12, and the estimated NIE is 0.069. Previous studies also support that these two CpG cites play important roles in tumor or lung cancer patients’ survival, which validates our findings. For example, the gene H19 was found to be related to the growth of lung cancer, and the methylation level of gene H19 was negatively correlated with tobacco smoking (Matouk et al., 2015; Bouwland-Both et al., 2015). Although cg24129177 is not located in any gene, Hair (2014) found that it was associated with body mass index (BMI), which was found to be related to the survival of lung cancer (Shepshelovich et al., 2019). As for cg07690349, Wakata et al. (2015) indicated that gene MUC5B where cg07690349 is located in is treated as a favourable prognostic biomarker for lung cancer patients with NSCLC carrying EGFR mutations. Furthermore, the combination of MUC5B and TTF-1 expression plays an important role in identifying adenocarcinomas from squamous cell carcinomas with prognostic significance in lung adenocarcinoma patients (Nagashio et al., 2015).

**Table 2:**
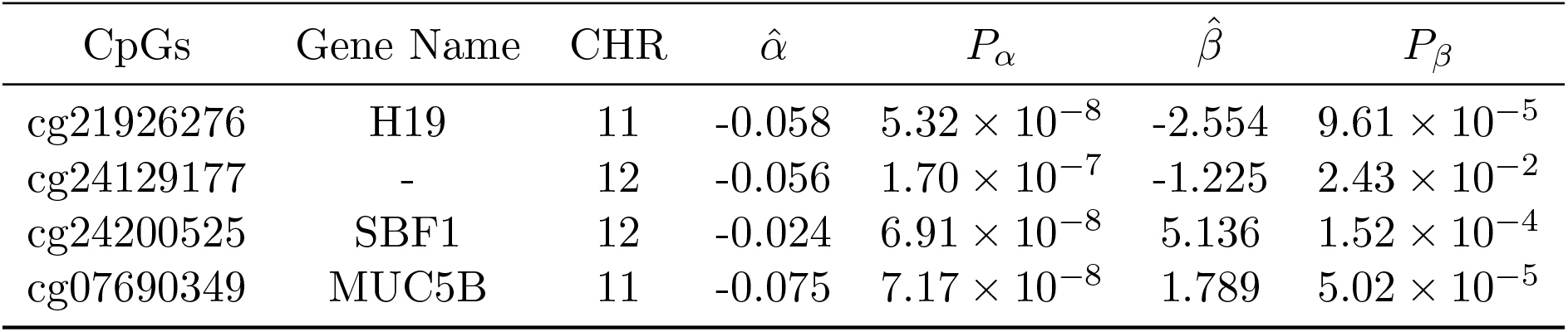
The identified DNA methylation CpG sites and estimated effects using CoxMKF with pre-specified *q*_1_ = 5 × 10^−4^, *q*_2_ = 0.1, *η* = 0.05 and *B* = 25. CHR stands for the chromosome.

| CpGs | Gene Name | CHR | $\hat{\alpha}$ | $P_\alpha$ | $\hat{\beta}$ | $P_\beta$ |
| --- | --- | --- | --- | --- | --- | --- |
| cg21926276 | H19 | 11 | -0.058 | $5.32 \times 10^{-8}$ | -2.554 | $9.61 \times 10^{-5}$ |
| cg24129177 | - | 12 | -0.056 | $1.70 \times 10^{-7}$ | -1.225 | $2.43 \times 10^{-2}$ |
| cg24200525 | SBF1 | 12 | -0.024 | $6.91 \times 10^{-8}$ | 5.136 | $1.52 \times 10^{-4}$ |
| cg07690349 | MUC5B | 11 | -0.075 | $7.17 \times 10^{-8}$ | 1.789 | $5.02 \times 10^{-5}$ |

We also apply HIMAsurvival to this data. In the preliminary screening of HIMAsurvial, one can choose to select the top mediators that are associated with either the exposure or the outcome. Since CoxMKF conducts the preliminary screening based on exposure-mediator associations, we also adopt the exposure-mediator associations for the preliminary screening procedure in HIMAsurvival for fair comparison. HIMAsurvival identifies five DNA methylation CpG sites: cg02178957, cg21926276, cg26387335, cg24200525 and cg07690349. Among these five DNA methylation CpG sites, the last four CpG sites were also identified in Luo et al. (2020). Besides, CoxMKF and HIMAsurvival identify 3 common DNA methylation CpG sites: cg21926276, cg24200525, and cg07690349. The result is consistent with our simulation studies that HIMAsurvival might have higher (maybe inflated) power to detect DNA methylation CpG sites but at the cost of inflated FDR in finite-sample settings.

As for the computational efficiency, the average running time of CoxMKF on this data set is 27.77 minutes with a server of Intel Xeon Silver 4116 CPU and 64 GB RAM memory, which is close to that of HIMAsurvival (26.63 minutes). Actually, the most time-consuming step is the screening process (25.89 minutes), which is a common step shared by both CoxMKF and HIMAsurvival. Notably, the multiple knockoffs generation and inference can be performed in parallel, which is no more costly than the single knockoff filter. Therefore, CoxMKF remains computationally efficient even when dealing with high-dimensional data.

In summary, we apply CoxMKF to the lung cancer data from the TCGA project and discover four DNA methylation CpG sites from the smoking status to the survival of lung cancer patients, of which most DNA methylation CpG sites are supported by previous studies. These four DNA methylation CpG sites might be helpful for researchers to design interventions to improve the treatment for lung cancer patients.

## 5 Discussion

In this paper, we propose a novel method CoxMKF for high-dimensional mediation analysis with a survival outcome. To achieve finite-sample FDR control, we incorporate a powerful procedure AKO for variable selection with a survival outcome. Simulation studies show that the proposed CoxMKF can achieve finite-sample FDR control compared to the competing method HIMAsurvival. We further identify four DNA methylation CpG sites in the pathway from smoking to lung cancer patients’ survival using the TCGA lung cancer data.

We highlight some advantages of CoxMKF here. First, the proposed CoxMKF applies AKO to conduct variable selection with finite-sample FDR control, which is more stable than the traditional model-X knockoffs method. Second, we propose two new feature statistics CCD and MSC for a survival outcome to extend the application of knockoff procedure to survival outcomes. Third, the proposed CoxMKF is computationally efficient even when dealing with high-dimensional data.

There are also some limitations of our proposed method. First, CoxMKF might be slightly conservative as it aims to provide a guarantee of FDR control in finite-sample settings. Second, we did not consider the correlation structure among multiple mediators. It might help increase power if the correlation structure is taken into account. In addition, it is also interesting to develop methods for other types of survival outcomes such as interval-censored outcomes.

## Supplementary Materials

**Table S1:**
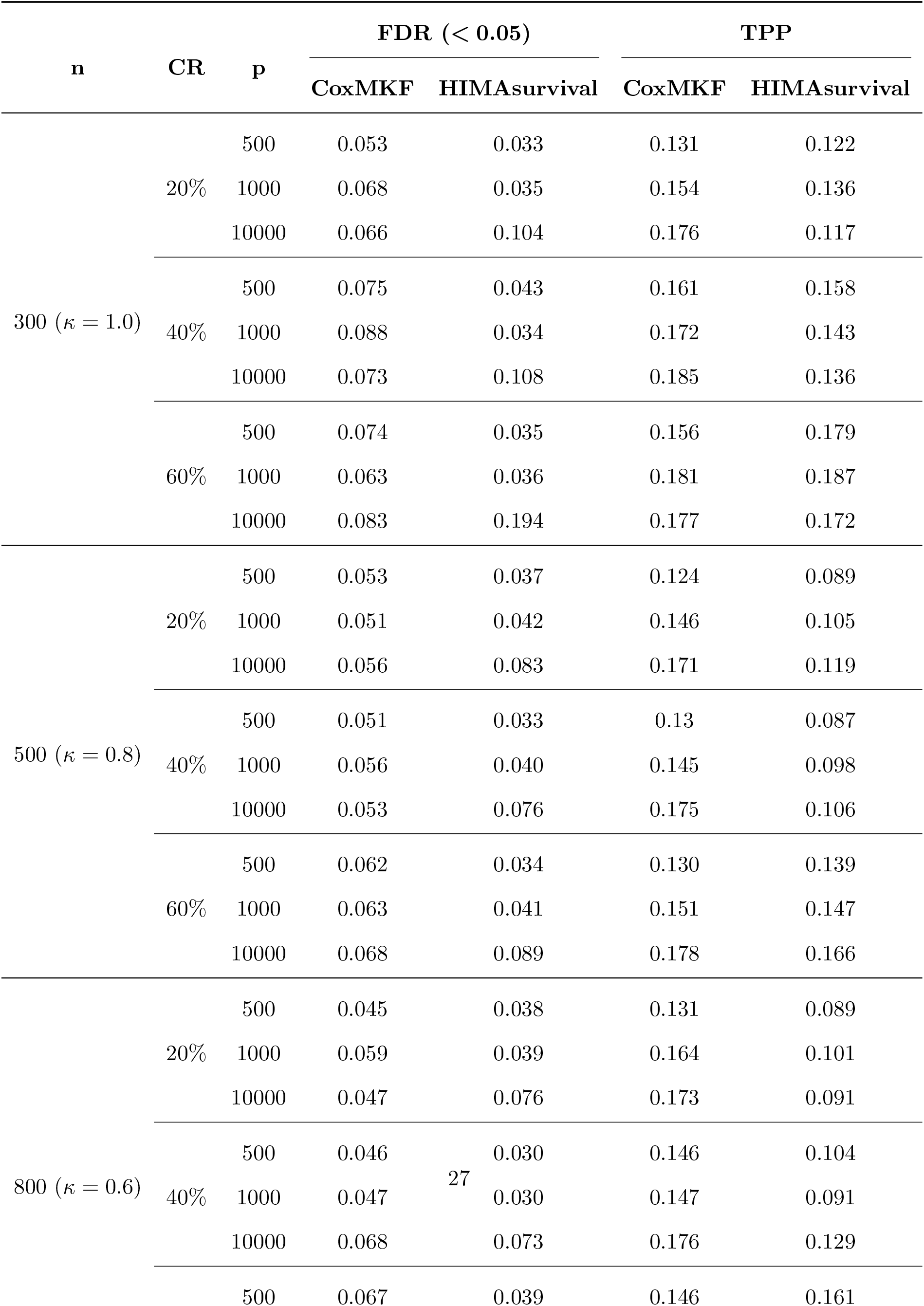
The standard errors of FDR and TPP of mediator selection across 500 simulations with *q*_1_ = 0.2, *q*_2_ = 0.05, *η* = 0.1 and the number of AKO’s bootstrap replications *B* = 25. The number of true mediators is 8. CR stands for the censoring rate. *κ* is a scaling factor to shrink the coefficients so that TPP is not too close to 1.

**Table S2:** The FDR and TPP of mediator selection across 500 simulations with *q*_1_ = 0.05, *q*_2_ = 0.02, *η* = 0.3 and the number of AKO’s bootstrap replications *B* = 50. We set *α_j_* = *κ* · 0.5 for *j* = 1, … , 12, *β_j_* = *κ* · 0.4 for *j* = 1, … , 10, *β*_13_ = *β*_14_ = *κ* · 0.5, and *κ* = 1, 0.8, 0.75 for *n* = 300, 500, 800, respectively. The rest elements of ***α*** and ***β*** are set to be 0. The number of true mediators is 10. CR stands for the censoring rate.

| n | CR | p | FDR ($< 0.02$) | | TPP | |
| --- | --- | --- | --- | --- | --- | --- |
|  |  |  | CoxMKF | HIMAsurvival | CoxMKF | HIMAsurvival |
| 300 ($\kappa = 1.00$) | 20% | 500 | 0.019 | 0.009 | 0.797 | 0.929 |
|  |  | 1000 | 0.016 | 0.011 | 0.738 | 0.926 |
|  |  | 10000 | 0.017 | 0.075 | 0.498 | 0.908 |
|  | 40% | 500 | 0.019 | 0.011 | 0.782 | 0.846 |
|  |  | 1000 | 0.018 | 0.014 | 0.713 | 0.839 |
|  |  | 10000 | 0.018 | 0.089 | 0.480 | 0.822 |
|  | 60% | 500 | 0.018 | 0.016 | 0.727 | 0.741 |
|  |  | 1000 | 0.019 | 0.023 | 0.665 | 0.747 |
|  |  | 10000 | 0.018 | 0.163 | 0.439 | 0.745 |
| 500 ($\kappa = 0.80$) | 20% | 500 | 0.014 | 0.012 | 0.848 | 0.953 |
|  |  | 1000 | 0.019 | 0.008 | 0.792 | 0.932 |
|  |  | 10000 | 0.010 | 0.053 | 0.586 | 0.916 |
|  | 40% | 500 | 0.020 | 0.003 | 0.813 | 0.784 |
|  |  | 1000 | 0.019 | 0.008 | 0.765 | 0.787 |
|  |  | 10000 | 0.016 | 0.051 | 0.543 | 0.770 |
|  | 60% | 500 | 0.014 | 0.001 | 0.808 | 0.793 |
|  |  | 1000 | 0.019 | 0.010 | 0.748 | 0.773 |
|  |  | 10000 | 0.013 | 0.288 | 0.471 | 0.663 |
| 800 ($\kappa = 0.75$) | 20% | 500 | 0.015 | 0.007 | 0.975 | 0.996 |
|  |  | 1000 | 0.011 | 0.013 | 0.950 | 0.997 |
|  |  | 10000 | 0.012 | 0.048 | 0.864 | 0.973 |
|  | 40% | 500 | 0.015 | 0.006 | 0.972 | 0.991 |
|  |  | 1000 | 0.020 | 0.009 | 0.920 | 0.970 |
|  |  | 10000 | 0.012 | 0.040 | 0.858 | 0.923 |
|  | 60% | 500 | 0.017 | 0.006 | 0.958 | 0.955 |
|  |  | 1000 | 0.015 | 0.010 | 0.950 | 0.959 |
|  |  | 10000 | 0.016 | 0.143 | 0.786 | 0.694 |

**Table S3:** The standard errors of FDR and TPP of mediator selection across 500 simulations with *q*_1_ = 0.05, *q*_2_ = 0.02, *η* = 0.3 and the number of AKO’s bootstrap replications *B* = 50. We set *α_j_* = *κ* · 0.5 for *j* = 1, … , 12, *β_j_* = *κ* · 0.4 for *j* = 1, … , 10, *β*_13_ = *β*_14_ = *κ* · 0.5, and *κ* = 1, 0.8, 0.75 for *n* = 300, 500, 800, respectively. The rest elements of ***α*** and ***β*** are set to be 0. The number of true mediators is 10. CR stands for the censoring rate.

| n | CR | p | FDR ($< 0.02$) | | TPP | |
| --- | --- | --- | --- | --- | --- | --- |
|  |  |  | CoxMKF | HIMAsurvival | CoxMKF | HIMAsurvival |
| 300 ($\kappa = 1.00$) | 20% | 500 | 0.045 | 0.029 | 0.145 | 0.099 |
|  |  | 1000 | 0.044 | 0.031 | 0.156 | 0.095 |
|  |  | 10000 | 0.057 | 0.088 | 0.185 | 0.112 |
|  | 40% | 500 | 0.047 | 0.034 | 0.158 | 0.147 |
|  |  | 1000 | 0.048 | 0.038 | 0.171 | 0.142 |
|  |  | 10000 | 0.056 | 0.116 | 0.190 | 0.153 |
|  | 60% | 500 | 0.046 | 0.048 | 0.177 | 0.189 |
|  |  | 1000 | 0.047 | 0.057 | 0.188 | 0.182 |
|  |  | 10000 | 0.060 | 0.223 | 0.201 | 0.189 |
| 500 ($\kappa = 0.80$) | 20% | 500 | 0.037 | 0.033 | 0.130 | 0.073 |
|  |  | 1000 | 0.047 | 0.029 | 0.147 | 0.100 |
|  |  | 10000 | 0.042 | 0.071 | 0.177 | 0.120 |
|  | 40% | 500 | 0.045 | 0.021 | 0.163 | 0.202 |
|  |  | 1000 | 0.045 | 0.033 | 0.162 | 0.204 |
|  |  | 10000 | 0.054 | 0.081 | 0.192 | 0.198 |
|  | 60% | 500 | 0.041 | 0.028 | 0.141 | 0.188 |
|  |  | 1000 | 0.047 | 0.032 | 0.161 | 0.180 |
|  |  | 10000 | 0.042 | 0.388 | 0.201 | 0.215 |
| 800 ($\kappa = 0.75$) | 20% | 500 | 0.035 | 0.026 | 0.049 | 0.021 |
|  |  | 1000 | 0.032 | 0.034 | 0.067 | 0.018 |
|  |  | 10000 | 0.064 | 0.076 | 0.181 | 0.097 |
|  | 40% | 500 | 0.036 | 0.026 | 0.054 | 0.038 |
|  |  | 1000 | 0.043 | 0.029 | 0.103 | 0.067 |
|  |  | 10000 | 0.036 | 0.063 | 0.124 | 0.113 |
|  | 60% | 500 | 0.037 | 0.022 | 0.064 | 0.080 |
|  |  | 1000 | 0.039 | 0.029 | 0.072 | 0.084 |
|  |  | 10000 | 0.041 | 0.286 | 0.160 | 0.229 |

## Supplementary Figures

**Figure S1:**
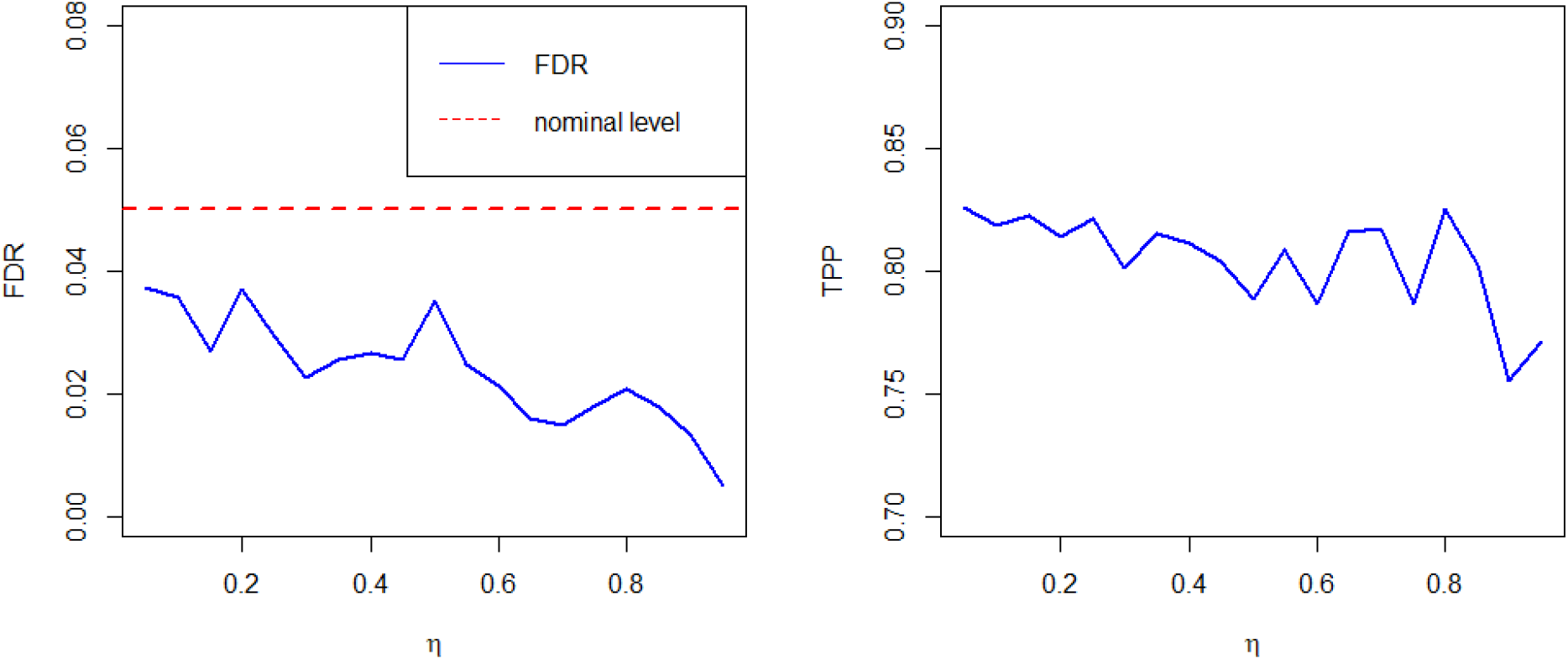
FDR and TPP of CoxMKF against different values of *η* across 500 simulations. The sample size is *n* = 300 and the number of candidate mediators is *p* = 10 000. The censoring rate is controlled at 40%. We fix the first ten elements of ***α*** to be (0.55, 0.45, −0.40, −0.45, 0.50, 0.60, −0.40, −0.46, −0.40, 0.50)^⊤^, the first twelve elements of ***β*** to be (0.52, 0.45, 0.40, 0.40, −0.54, −0.60, −0.40, −0.50, 0, 0, 0.40, −0.80)^⊤^, and the rest elements of ***α*** and ***β*** are set to be 0. We set *q*_1_ = 0.2, *q*_2_ = 0.05 and *B* = 25.

